# Magnetomyography: A novel modality for non-invasive muscle sensing

**DOI:** 10.1101/2024.04.15.588623

**Authors:** Richy Yun, Gabriel Gonzalez, Isabel Gerrard, Richard Csaky, Debadatta Dash, Evan Kittle, Nishita Deka, Dominic Labanowski

## Abstract

The measurement of magnetic fields generated by skeletal muscle activity, called magnetomyography (MMG), has seen renewed interest from the academic community in recent years. Although studies have demonstrated complex models of MMG and experiments classifying between different movements using MMG, there has yet to be time frequency analysis of MMG as well as concurrent recordings of MMG and its electrical counterpart, surface electromyography (sEMG). Here, we aim to better understand MMG in the context of sEMG by simultaneously recording both modalities during various muscle contraction tasks. We found that, similar to sEMG, MMG shows highly linearly correlated power to the degree of muscle contraction, has a unimodal distribution in spectral power, and can detect changes in muscle fatigue via changes in the spectral distribution. One main difference we found was that MMG typically has more high frequency content compared to sEMG, even when accounting for the filtering induced by the size of the sEMG electrodes. We additionally demonstrate empirically the decrease in MMG power due to distance from the arm and show MMG decreases slower than the inverse square law and can be measured up to 50 mm from the surface of the skin. Finally, we were able to capture MMG with non-OPM sensors showing that sensor technology has made great strides towards enabling MMG applications.

## Introduction

Measurement of magnetic fields produced by the human body, or biomagnetism, was first demonstrated with recordings of the heart, or magnetocardiography (MCG), by Baule and McFee using a superconducting quantum inference device (SQUID) (Baule & McFee, 1963). The first magnetic recordings of the brain and muscle, magnetoencephalography (MEG) and magnetomyography (MMG) respectively, were subsequently demonstrated by Cohen (Cohen, 1968; Cohen & Givler, 1972). Both MCG and MEG were continuously developed over the decades and are used today, albeit seldomly. However, MMG has been understudied because existing magnetometers with sufficient sensitivities are difficult to implement, making its electrical counterpart, surface electromyography (sEMG), the dominant technology for most applications.

Recently, however, there have been major advances in magnetic sensors – including the miniaturization of optically pumped magnetometers (OPMs) (Shah & Wakai, 2013), high performance tunneling magnetoresistance (TMR) sensors (Zuo et al., 2020), and novel sensor technologies like acoustically driven ferromagnetic resonance (ADFMR) sensors (Labanowski et al., 2016). These sensors have sensitivities high enough to capture MMG while being mobile and modular, providing numerous benefits for assessing MMG compared to SQUIDs.

Several studies have demonstrated robust recordings of MMG and have applied analyses including stimulus triggered averages of muscle activity (Broser et al., 2018), decoding of individual motor units (Broser et al., 2023), and even discrimination between movements that rival results obtained with sEMG (Greco et al., 2023). Models and simulations have also shown MMG has higher specificity and better localization compared to sEMG, furthering excitement for the field (Arekhloo et al., 2022; Klotz et al., 2023).

Despite the increase in the number of studies on MMG, literature typically reports filtered time traces or stimulus triggered averages of MMG. Previous reports have even compared the accuracy of gesture discrimination between MMG and sEMG (Greco et al., 2023), but there has yet to be a comprehensive time-frequency characterization of voluntary MMG or an empirical comparison between MMG and sEMG. Though the origin of both modalities is the currents traveling through muscle fibers, there are notable differences between how they propagate. sEMG signals are warped by the conductive properties of the signal path – tissue, skin, and the impedance at the sensor interface – whereas the body is permeable to magnetic fields leaving MMG signals unaffected (Arekhloo et al., 2022; Hansen et al., 2010). This distortion introduced to sEMG is highly nonlinear and depends heavily on the subject and day-to-day environmental conditions, introducing difficulties in generalization and scaling with sEMG. As such, we need a better understanding of not only the characteristics of MMG but also its differences from sEMG.

To that end, we simultaneously captured MMG and bipolar sEMG. This study is the first to show characterization of MMG compared to sEMG, including time frequency analysis with respect to muscle contraction and muscle fatigue, how MMG falls off as a function of distance from the signal source, and voluntary MMG captured with various sensors. We believe the results provide context for better understanding previous findings, lay the groundwork for future MMG studies, and demonstrate MMG recordings with different sensors to highlight the applicability of MMG.

## Methods

### Data collection

#### Sensors and acquisition

We used three different magnetometers for our measurements: QZFM Gen-2 (QuSpin, Inc.), Aichi DJ (Aichi Steel Corporation), and TDK Nivio (TDK Corporation) (Figure 1a). Table 1 shows various metrics for each sensor. The sensors were strapped to subjects using a combination of custom 3D printed holders and Velcro straps.

**Figure 1.**
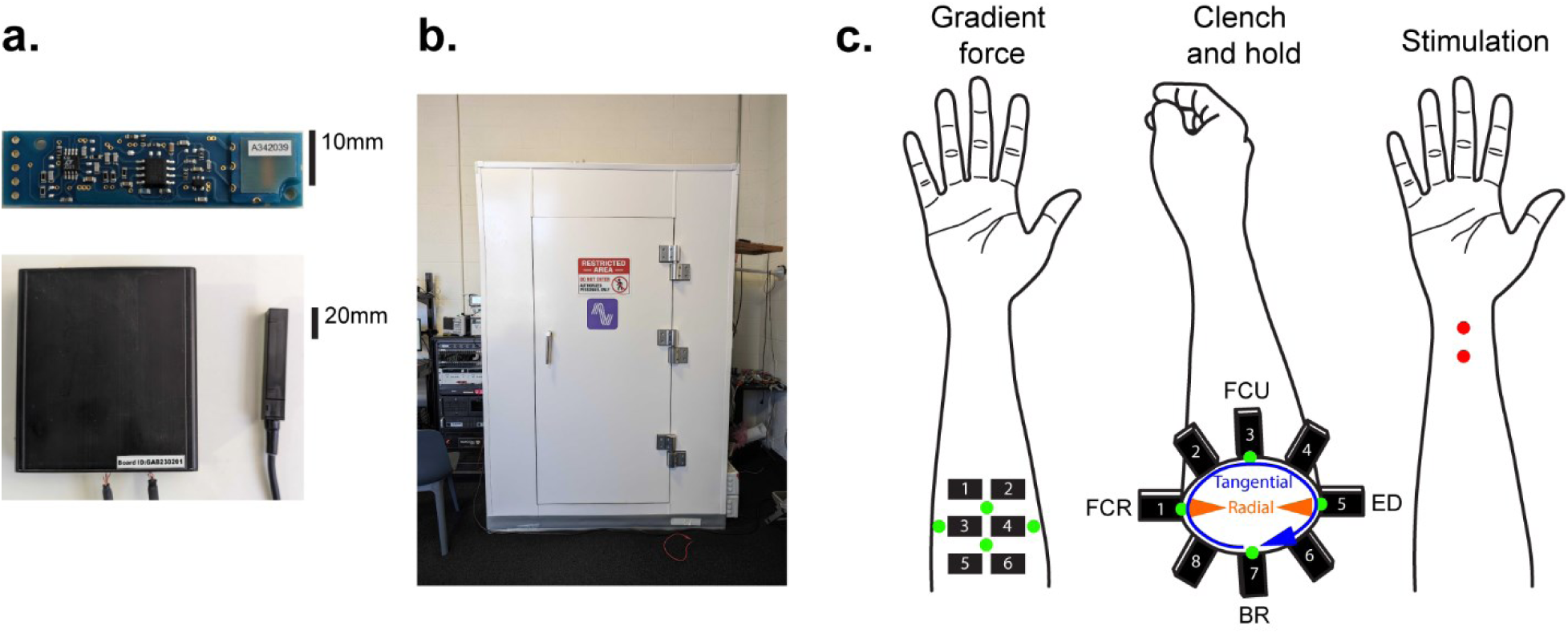
Experimental setup. **a,** non-OPM sensors used in this study are the Aichi DJ (top) and TDK Nivio (bottom, readout electronics on the left and sensor body on the right). **b,** Magnetically shielded room used for all experiments. **c,** OPM and sEMG sensor positions for the gradient force task (left); only the radial axis was recorded for each OPM sensor. Sensor positions for clench and hold task (middle) with labeled sensors roughly corresponding to flexor carpi radialis (FCR), flexor carpi ulnaris (FCU), extensor digitorum (ED), and brachioradialis (BR). Both radial and tangential axes were recorded for each OPM sensor. Green dots for both denote the center of sEMG pairs. Finally, we also delivered stimulation to activate the median nerve using bipolar sEMG electrodes (right). Red dots denote locations of each sEMG electrode.

**Table 1.** Comparison of different magnetometers used in this study.

|  | QuSpin QZFM Gen-2 | Aichi DJ | TDK Nivio |
| --- | --- | --- | --- |
| **Technology** | OPM | MI | xMR |
| **Noise at 100Hz** | 10 fT/$\sqrt{\text{Hz}}$ | 2 pT/$\sqrt{\text{Hz}}$ | 500 fT/$\sqrt{\text{Hz}}$ |
| **Bandwidth** | 0.1 – 200 Hz | 0.1 – 10 kHz | 0.1 – 40 kHz |
| **Operating range** | 5 nT | 1 $\mu\text{T}$ | 45 $\mu\text{T}$ |
| **Number of axes** | 2 orthogonal | 1 | 1 |
| **Sensor volume** | 26.4×16.6×12.4 mm | 55×13.5×5 mm | 70×12×12 mm |
| **Readout electronics volume** | 3.1×11.0×16.5 cm | N/A | 12×10×3 cm |

All sEMG recordings were performed with single-use, bipolar, MRI safe, pre-gelled sEMG electrodes with non-ferrous contacts (EL508, Biopac Systems Inc.). The gelled area was roughly 10 mm in diameter, and the center-to-center inter-electrode spacing of bipolar pairs was kept to 20 mm. The sEMG leads were also non-ferrous to limit introduction of artifactual magnetic fields (LEAD108C, Biopac Systems Inc.). Comparisons with standard sEMG electrodes and leads showed no differences in recorded data.

To measure the grip strength of subjects we used a pneumatic dynamometer bulb attached to an analog pressure sensor (TR1-0030A-101, Merit Sensor). To measure force gradient produced by finger flexion we used a FlexiForce A201 (Tekscan) and custom readout electronics.

All sensors provided analog signals which were digitized using NIDAQ ADC modules (National Instruments, 16-bit accuracy for OPMs, 24-bit accuracy for all other sensors) and collected with custom Python data acquisition code.

#### Magnetically shielded room

We used a MuROOM (Magnetic Shield Corporation) with inside dimensions of 1.3×1.3×2 meters to prevent sensor saturation and contamination with ambient noise sources (Figure 1b). The shielded room had around 25,000 fold attenuation of residual fields at DC and up to 8000 fold attenuation at AC. The room was degaussed prior to each recording to guarantee maximum rejection of ambient fields. All recordings were performed inside the shielded room.

#### Participants and tasks

We had a total of 5 subjects participate in the study (4 males and 1 female in their 20’s and 30’s, all right-handed). The subjects, all employees of Sonera Inc., gave their informed consent to participate in each experiment.

Our study employed three different tasks. The first was a gradient force task in which the subject rested the ventral forearm flush on an acrylic table and slowly increased downward pressure onto the pressure sensor using their middle finger up to 100% maximum voluntary contraction (MVC) across 5 seconds before slowly ramping back down to relax. This task was cued with an audio tone. 2 subjects participated in this task.

The second involved the clench and hold task in which the subject was instructed to clench their fist at 100% maximum voluntary contraction (MVC) and hold for 10 seconds. Each trial was separated by 5 seconds of rest in which the subjects fully relaxed their arm. The start of the contraction and relaxation periods were cued by an audio tone, and subjects continued the task until they could not hold the contraction for the full 10 seconds. 5 subjects participated in this task when using a wristband of 16 dual-axis OPMs and four bipolar sEMG channels (Figure 1c). 2 subjects participated in this task when using OPM sensors at different distances, TDK sensors, and Aichi sensors.

Lastly, we used the same sEMG electrodes and leads as recordings to deliver stimulation to the median nerve on one subject. A pair of electrodes were placed along the length of the arm on the midline of the ventral side of the wrist with 20 mm inter-electrode distance to activate the median nerve (Pease et al., 2007). A Digitimer DS7A was used to deliver stimulation and was triggered by an Arduino microcontroller. The stimulus was monophasic and single pulse with 500 µs pulse width and alternating polarity each stimulus. The amplitude was adjusted in 1 mA increments until we observed roughly 80% of maximum recruitment as measured by sEMG response amplitudes (typically ∼30 mA). We then delivered the stimulation every 2 seconds with a jitter of 500 ms for 5 minutes.

#### Sensor placements

For the clench and hold task 8 OPM sensors and 4 bipolar sEMG sensor pairs were placed around the dominant forearm. To capture the largest muscle signals, we placed the sensors at 1/3 of the full forearm length from the antecubital fossa (crook of the elbow). The first OPM sensor was placed roughly over the flexor carpi radialis and each subsequent sensor was placed 45 degrees clockwise (Figure 1c). The first sEMG sensor was also placed over the flexor carpi radialis and each subsequent sensor was placed 90 degrees clockwise. The center of the bipolar sEMG electrodes were positioned to be directly below the corresponding OPM sensors.

For the gradient force task 6 OPM sensors were placed in a 3×2 grid on the other side of the acrylic surface in a custom 3D printed mount (∼2.5 cm distance from the surface of the skin).

For measurements with TDK and Aichi sensors as well as when using multiple OPM sensors at different distances the sensors were placed above the extensor digitorum and held by a custom 3D printed holder.

### Data analysis

#### Preprocessing

All data were captured at a 2 kHz sampling rate. Signals were first filtered at 60 Hz harmonics with a 2^nd^ order IIR notch filter with a q-factor of 30 to remove line noise. For analyses requiring further processing, we applied a band pass filter from 20 to 300 Hz with a 5^th^ order Butterworth filter.

To account for trial-to-trial variability, we aligned each trial to the time of maximum grip strength as subjects typically started with the strongest grip that loosened with time as muscles were fatigued (Supplementary Figure 1).

#### Median and mean frequencies

The median frequency (Stulen & De Luca, 1981) is defined as the frequency at which the sum of the spectrum is equal on both sides of the frequency:

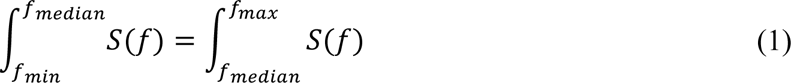

where *f*_*min*_ and *f*_*max*_ are the minimum and maximum frequencies of the bandwidth and *S*(*f*) is the spectrum of the signal.

The mean frequency (Stulen & De Luca, 1981) is defined as the average of frequencies weighted by the spectrum:

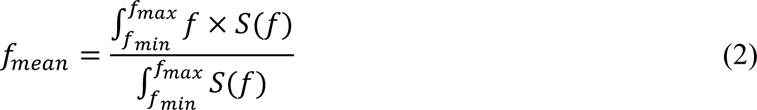

Both median and mean frequencies were calculated with a bandwidth of 20-300 Hz.

#### Calculation of distance dependence

The decrease in signal strength with respect to distance between the sensor and the signal source can be written as:

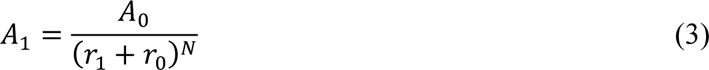

Where *A*_0_ and *A*_1_ are signal amplitudes at the source and sensor respectively, *r*_0_ is the distance from the source to the surface of the skin, *r*_1_is the distance from the surface of the skin to the sensor, and *N* is the rate of decrease in the signal.

However, the signal amplitude at the source and the distance between the source and the surface of the skin are both unknown. As such, there needs to be a second sensor at a different distance, which gives:

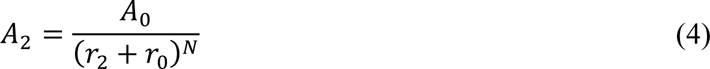

Dividing Equation 3 by Equation 4 and solving for *N* gives:

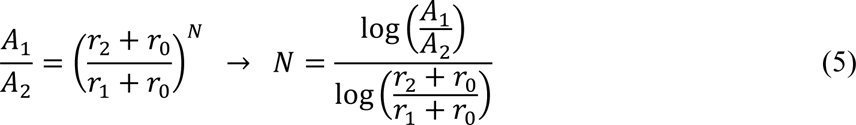

Equation 5 provides a method with which to calculate *N* as a function of *r*_0_. With three sensors at three distances Equation 5 can be modified to directly calculate *N*:

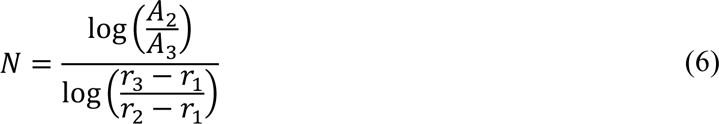

Note the subtraction of *r*_1_ in Equation 6 rather than addition as in Equation 5 as the distance between the sensors must be calculated.

#### Statistical analysis

All statistical analyses assumed non-parametric distributions of data and typically consisted of the one- or two-sided Wilcoxon signed rank (`scipy.stats.wilcoxon`) or rank sum (`scipy.stats.ranksums`) tests. Linear correlations were calculated using the Pearson correlation coefficient (`scipy.stats.pearsonr`). Statistical significance was determined at an alpha level of 0.05. Each analysis specifies the tests used to obtain the results.

## Results

### MMG magnitude is correlated with force of muscle contraction

The amplitude of sEMG has been shown to be directly correlated with the strength of muscle contraction, allowing it to be employed in applications in which varying contraction is a significant factor (Clancy et al., 2001; Disselhorst-Klug et al., 2009). To determine whether MMG also has the same correlation, we asked the subjects to press down with a finger with increasing strength over time and measured the flexor muscles on the forearm.

We calculated Pearson correlation coefficients between the measured force and sEMG signals and found highly linearly correlated changes in sEMG signal power with increase in force produced by finger flexion (Figure 2a), similar to previous literature (Disselhorst-Klug et al., 2009; Hof, 1997). We found the correlation to be consistent across the bandwidth with the Pearson correlation having a standard deviation of 0.02 (Figure 2a, right). Though there are reports of frequencies being unchanged or even having negative correlations with contraction strength, these results are often subject-specific, indicating they are due to variability in volume conductor properties and muscle composition (Moritani & Muro, 1987; Rodriguez-Falces et al., 2015).

**Figure 2.**
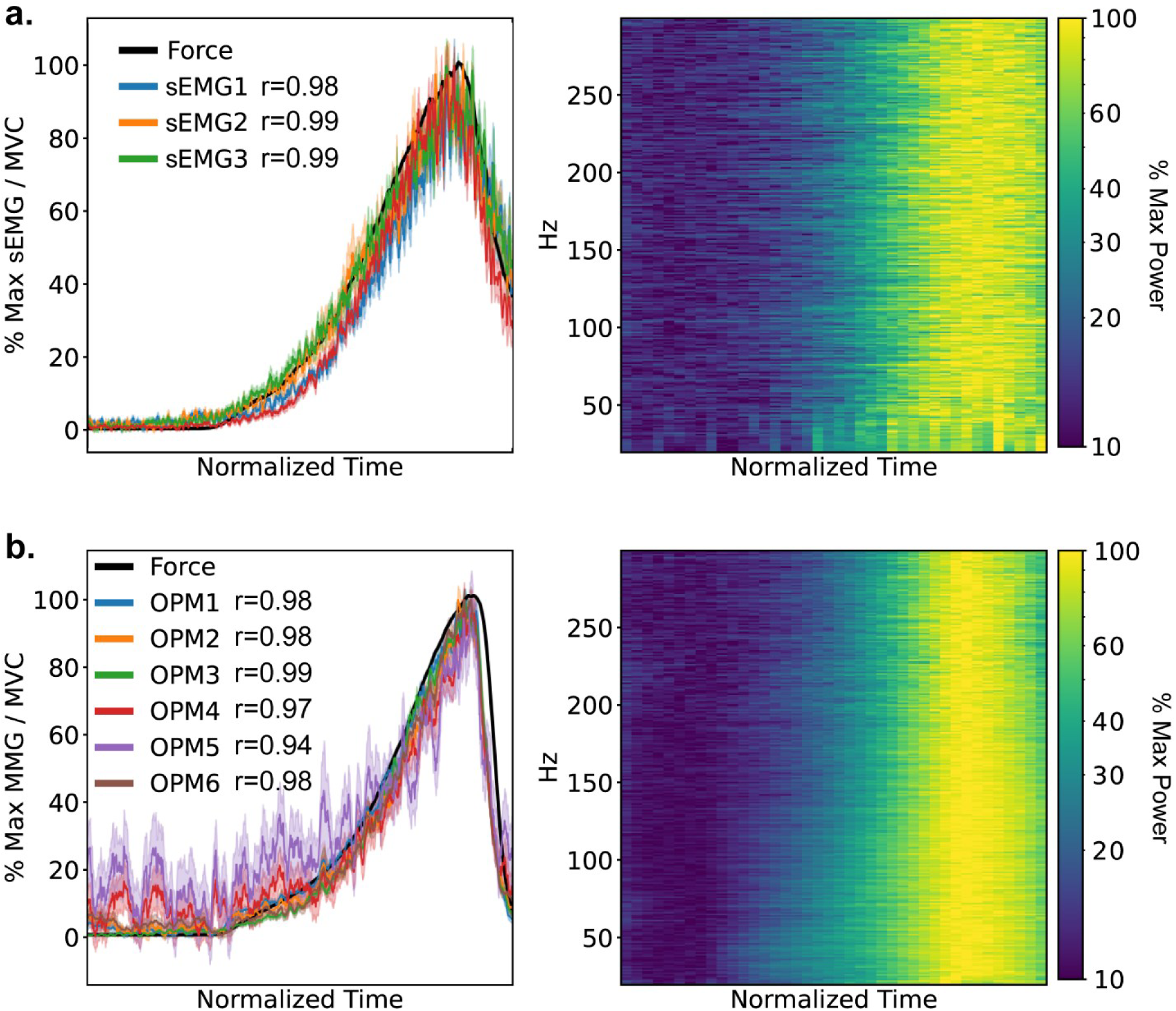
Gradient force of contraction and changes in sEMG and MMG signal strength. **a,** Total sEMG power between 20 and 300 Hz normalized to maximum power averaged across all trials as well as averaged measured force (left). As there was variability in timing of force between trials, we normalized the time from 10% to 100% of MVC to be uniform duration across trials. Pearson r correlation coefficients are shown for each channel compare with the measured force. The spectrogram of the sEMG signals normalized for each frequency for channel sEMG1 is shown on the right. **b,** Equivalent analysis applied to MMG signals, OPM1 is shown on the right.

Repeating the analyses on MMG showed similar results (Figure 2b). The correlation in MMG was similarly consistent across the bandwidth with the Pearson correlation also having a standard deviation of 0.02 (Figure 2b, right).

An additional result of note is the variability of MMG. Figure 2b left shows normalized power, but it is clear there are specific channels which show high responses to muscle contraction compared to others (i.e. high variability at low contraction strength suggests lower signal-to-noise ratio). sEMG channels that were similarly distanced all showed very similar signals across all channels. This potentially suggests higher spatial specificity of MMG, though additional investigation is necessary to draw firm conclusions.

Finally, we also assessed what percentage of maximum voluntary contraction (MVC) could be detected with either modality by capturing the average and standard deviation of the signal at rest. Setting a threshold to the average plus two times the standard deviation, we found MMG to be able to detect contraction strengths as small as sEMG (Supplementary Figure 2). We also found that the variance between the MMG channels was due to the placement of the sensors, with channels in high proximity having more similar detection thresholds.

### MMG has large bandwidth with relatively higher power at higher frequencies

After confirming that MMG power is correlated with the strength of contraction, we sought to compare the spectral distribution between MMG and sEMG. MMG frequency content was reported during its initial discovery (Cohen & Givler, 1972), but the recording device was roughly 4 cm from the surface of the skin and the measurements were of the upper arm and the palm. To expand upon these results, we recorded MMG and sEMG from around the forearm, as gesture recognition from a wearable placed on the wrist or the forearm is a major emerging application of neuromuscular recordings. The subjects were instructed to clench their fists at 100% MVC and sustain the muscle contraction for 10 seconds with 5 seconds of relaxation periods in between each trial.

Figure 3a shows the averaged force across a single session of the clench and hold task and the averaged spectrogram of sEMG and MMG measured at the extensor digitorum for a single subject. The timing of the two signals aligns well with the exerted force, and both modalities show significant power up to 100s of Hz. We were able to verify MMG signal power at 300 Hz even though the frequency response of the OPM sensors we used start decreasing at 100 Hz and roughly -3.5 dB at 300 Hz (Supplementary Figure 3), signifying MMG may have strong high frequency content.

**Figure 3.**
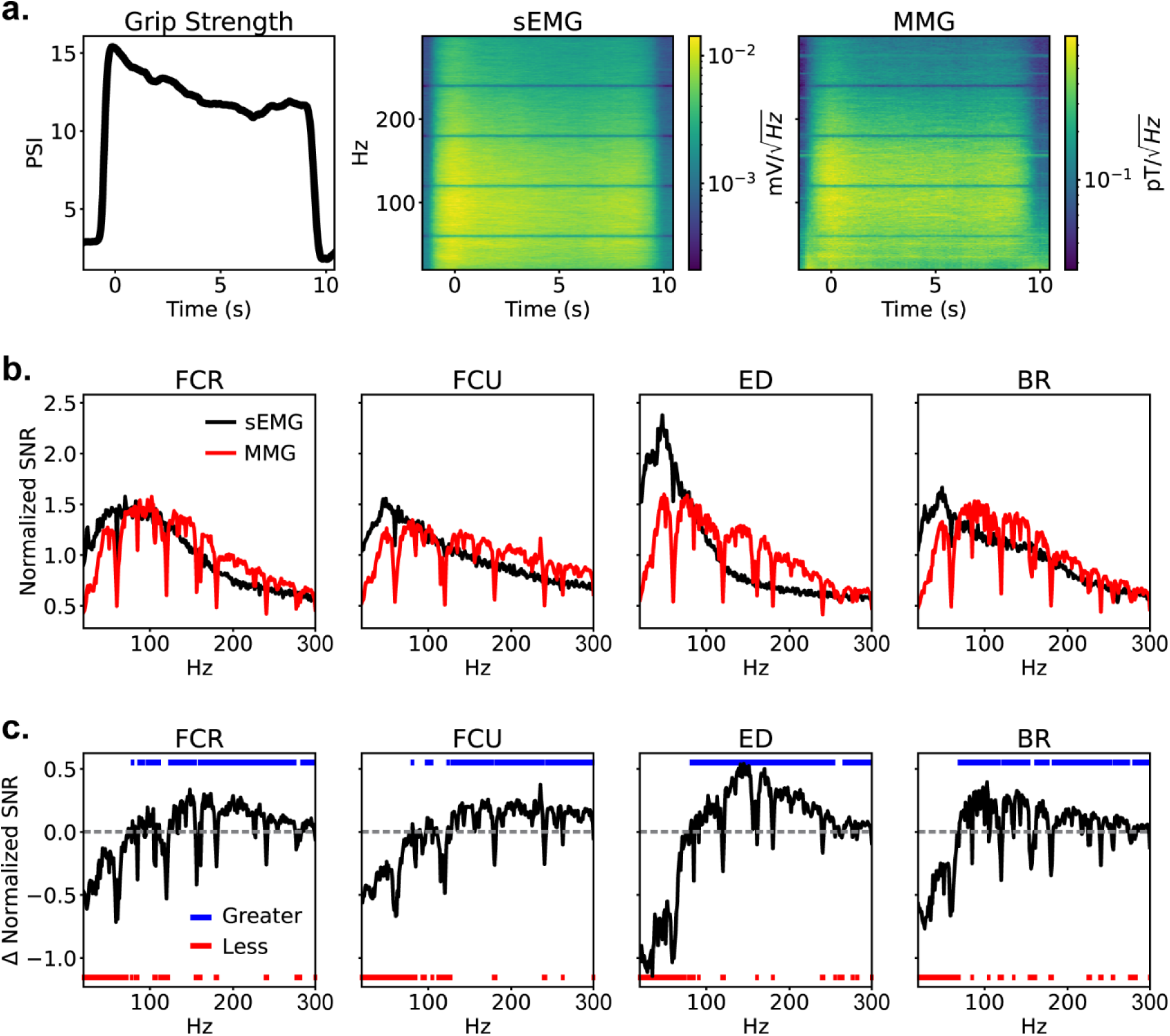
Frequency analysis of sEMG and MMG during the clench and hold task. **a,** Example grip strength, single channel sEMG spectrogram, and single channel MMG spectrogram averaged across an entire session. **b,** Signal to noise ratio (SNR) of spectra during muscle contraction compared to during rest for each sEMG channel and corresponding MMG channel. The SNR was normalized by dividing by the average SNR within the bandwidth to properly compare the frequency specific SNR between modalities. **c,** Difference in normalized SNR between MMG and sEMG (MMG SNR – sEMG SNR). The horizontal dashed line is at 0, or when MMG and sEMG have equal SNR. The colored dots show where MMG SNR is greater than (blue) or less than (red) sEMG SNR (Wilcoxon signed rank test). Note the cases in which MMG SNR is less than sEMG SNR at higher frequencies typically fall within 60 Hz harmonics as the OPM sensors used had greater line noise or at frequencies higher than 200 Hz likely due to the frequency response of the OPM sensors (Supplementary Figure 3). See Supplementary Figure 4 for MMG SNR corrected to account for the frequency response.

To compare between the two modalities, we calculated the signal-to-noise ratio (SNR) for each trial by capturing the spectra during contraction and dividing by the average spectra of when the subject is at rest. The SNR for each trial was subsequently normalized to emphasize relative power between frequencies by dividing by the average power within a 20-300 Hz bandwidth, then averaged across all sessions and subjects (Figure 3b). Note that both axes of the OPM sensors (radial and tangential) were used.

We found that MMG typically had higher relative power in the higher frequencies compared to sEMG on all directly comparable sensor pairs. By calculating the difference between the normalized SNR of MMG and sEMG, MMG has significantly greater relative power at frequencies greater than 100 Hz and up to 300 Hz even without accounting for the frequency response of the OPM sensors (Figure 3c). This suggests MMG may provide more reliable recordings of motor unit action potentials as well as more precise localization of active muscle fibers compared to sEMG, though we were unable to test these hypotheses due to limitations of commercially available sensors.

To have the comparison be as equivalent as possible, we also corrected the MMG spectra with the OPM frequency response. The sEMG signals are also low pass filtered, mainly due to the size of the electrodes. As a result, we estimated the change in frequency response by using previously reported spatial frequency response and assuming a conduction velocity of 4 m/s (Merletti & Muceli, 2019) to match that of a 3 mm wide electrode as the OPM vapor cell is a 3 mm wide cube. The resulting normalized SNR shows a slight reduction in the difference at high frequencies and pushes the difference up to roughly 150 Hz for FCR and FCU but maintains that MMG has significantly greater relative power at higher frequencies (Supplementary Figure 4).

### MMG can detect muscle fatigue

We assessed muscle fatigue by evaluating the temporal changes in MMG signal in the clench and hold task. The task was designed such that it continued until the subjects could not complete a full trial reflecting muscle fatigue. Though numerous methods of tracking muscle fatigue with sEMG signals exist (Rampichini et al., 2020), we focused on the decrease in average frequency (i.e. relative decrease in high frequency power) due to its spectral relevance (Bigland-Ritchie, 1981; Sometti et al., 2021).

To that end, we calculated the spectrogram of the signals for each channel in each trial. To determine changes in spectral content within trials (i.e. throughout the duration of holding the contraction) and across trials (i.e. throughout the duration of the session) we normalized the spectrograms either across time within trials or across trials (Figure 4, top).

**Figure 4.**
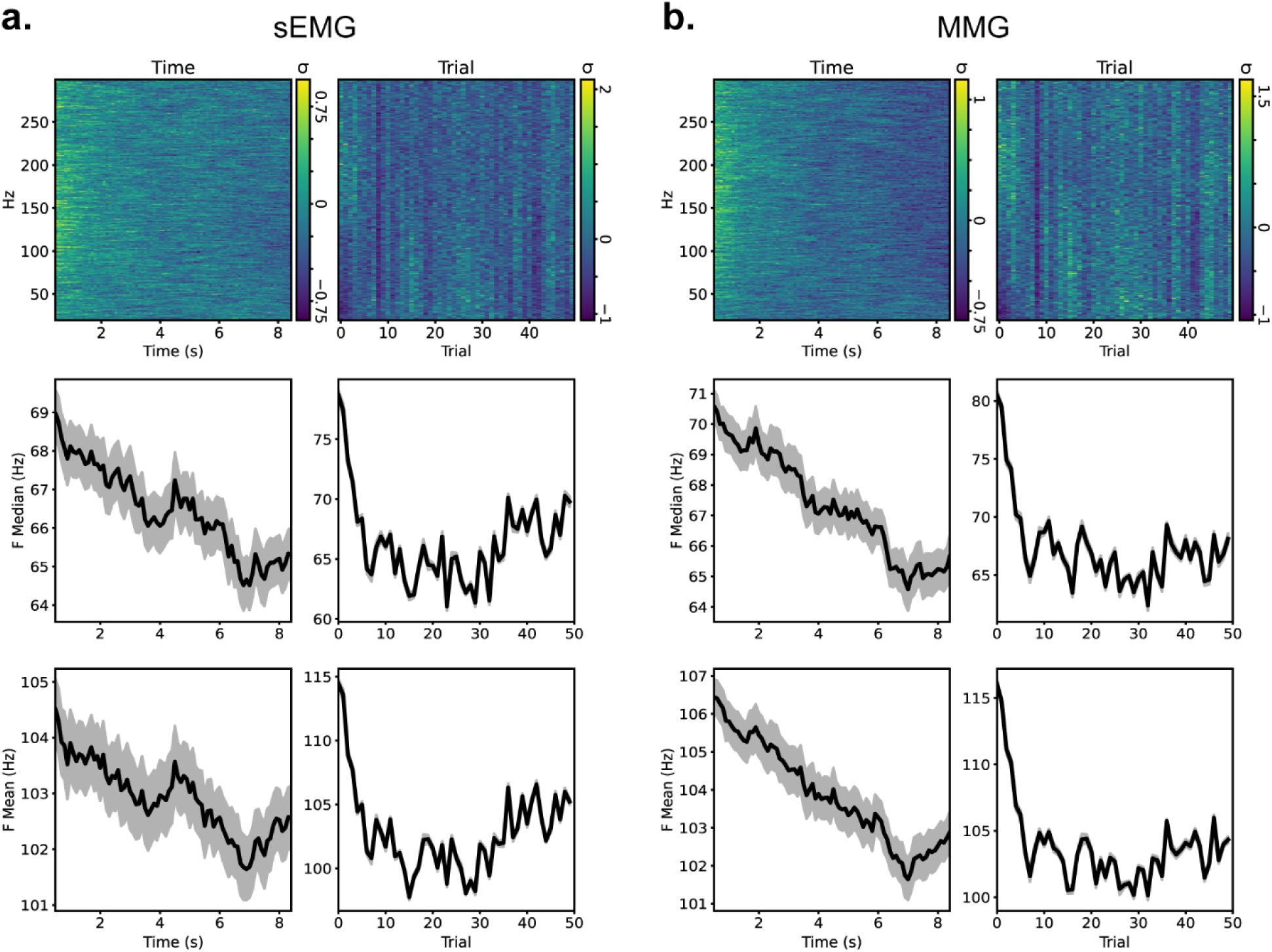
Changes in sEMG and MMG due to muscle fatigue. **a,** Example normalized spectrogram of sEMG positioned over the brachioradialis averaged for within trials (top left) and across trials (top right) for a single session. Median (middle) and mean (bottom) frequencies within and across trials. Shaded region shows standard error of the mean. **b,** Same analyses applied to radial MMG signals positioned at the same location.

From the spectrograms it was clear that most frequencies decreased over time but different frequencies behaved differently depending on the degree of fatigue (i.e. time within trials and number of trials within a session). As a result, we calculated the median and mean frequencies for both sEMG and MMG signals both within and across trials (Figure 4, middle and bottom).

There is a clear decrease in both measures both within and across trials showing that high frequency signals were decreasing faster, similar to previously reported results (Rampichini et al., 2020; Sometti et al., 2021).

Both median and mean frequencies reflected the same results, demonstrating lack of outlier influence. Interestingly, both measures decreased linearly within trials but had a large decrease within the first 10 trials before stabilizing across trials, or even going up in the later trials as in the case of sEMG in the example. We also found that MMG typically had larger change in the measures, as demonstrated in Figure 4.

Not all channels of sEMG and MMG displayed such changes. We assessed whether fatigue was detected by statistical comparison of the first second or first 10 trials compared with the last second or the last 10 trials. Fatigue was only noted if the difference was significant both within and across trials (Supplementary Figure 5). As each subject had slightly differing strategies of grasping the pressure bulb (position and angle of bulb in the hand, positions of the fingers around the bulb, etc.) as well as varying muscle structure, we observed subject dependent differences in which channels displayed fatigue. Note, however, that sEMG and MMG channels located in proximity typically showed similar changes.

### MMG signal strength as a function of distance typically decreases slower than the inverse square law

The magnetic field of a dipole falls off as a cube of the distance from the source, the field from a finite wire as the square of the distance, and the field from an infinite wire as the inverse of the distance. Although muscle fibers are often modeled as finite wires, the drop-off is typically considered to be the cube of the distance when recording very close to the source to become the inverse of the distance when further from the source (Arekhloo et al., 2023; Wijesinghe, 1991).

To empirically quantify the rate at which the signal decreases, we measured MMG signals over the extensor digitorum during a clench and release task for 20 trials across 2 subjects. MMG was captured with three OPM sensors measuring radially with 10 mm between the sensors, translating to vapor cells (i.e. the sensitive element of the sensors) at distances of 6.2, 28.6, and 51 mm from the skin (Figure 5a and b). The OPM sensors apply an internal bias field to cancel out external DC fields which can interact between sensors in close proximity resulting in sensor saturation, and greater than 10 mm distance between the sensor bodies often led to no signal captured in the furthest sensor. As a result, we could not take measurements at different distances.

**Figure 5.**
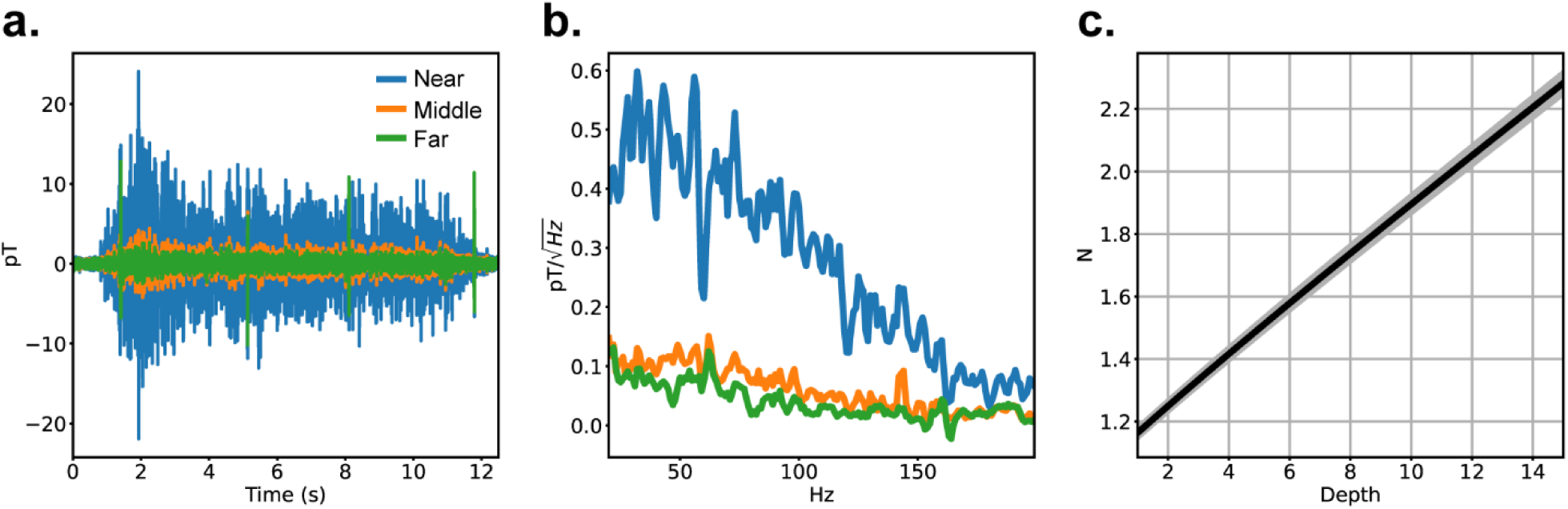
MMG and distance. **a,** Example filtered time domain signals from 3 OPM sensors at different distances during a single trial of clench and hold. **b,** Averaged spectra for each sensor. **c,** Rate of signal decrease with respect to distance (N) at hypothetical source depths. Shaded region shows standard error of the mean.

Using the measurements from the near and middle sensors, we can estimate the rate of decrease in the signal at the skin surface with respect to the hypothetical depth of the signal source (Figure 5c). In the forearm, the surface of the muscle is unlikely to be further than 10 mm from the surface of the skin (Gibney et al., 2010), meaning that the decrease of signal power is likely to be slower than the inverse square root law (N<2) at the skin surface. Using all three sensors we calculated the rate of decrease between the middle and far sensors to be 0.90 ± 0.05 (average ± SEM), showing the decrease quickly becomes linear.

### MMG can be captured with commercial non-OPM magnetometers

Although SQUIDs and OPMs have typically been used to detect MMG due to the high sensitivity requirements, we tested whether we could detect MMG with a commercial magnetoimpedance (MI) sensor, the Aichi DJ. Though previous studies have demonstrated MMG with non-OPM sensors, they were typically performed with custom hardware in research settings (Gielen et al., 1986; Zuo et al., 2020). To that end we took inspiration from Broser et al. and measured the abductor pollicis brevis while delivering stimulation to the median nerve at the wrist (Broser et al., 2018).

Three sessions on one subject were captured, each with a different type of sensor. The triggered averages of each are shown in Figure 6. It shows that the responses captured with the Aichi DJ is very similar to the response measured with sEMG (biphasic response from 5 – 15 ms). Signals recorded with OPMs resulted in a similar response but were not able to properly capture the sharp features. This may be due to the bandwidth limitations of the OPM (Supplementary Figure 3).

**Figure 6.**
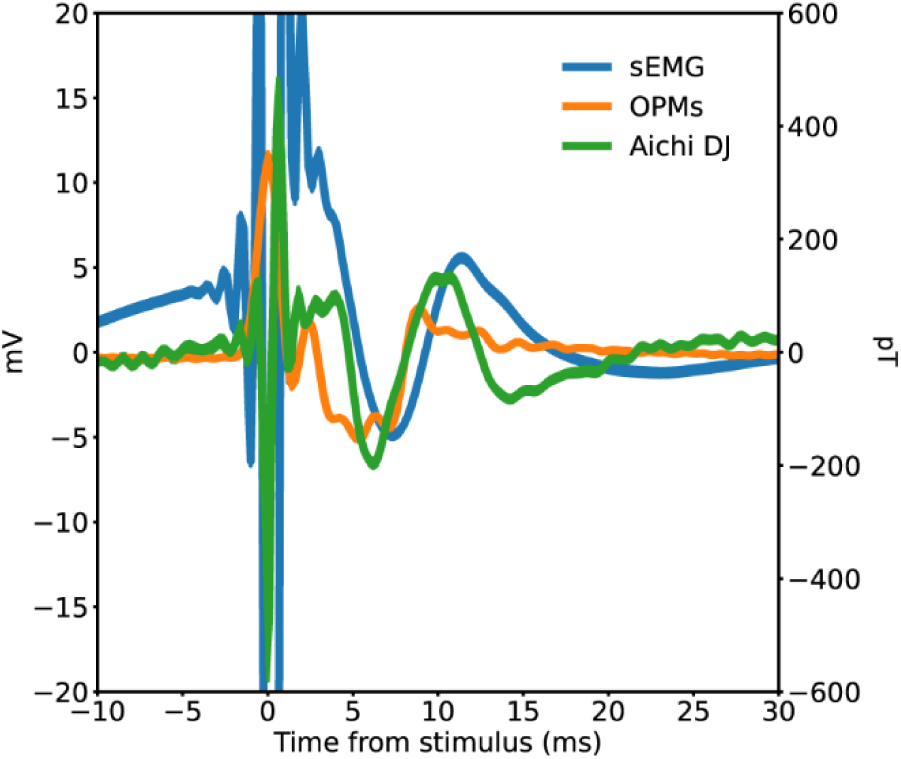
Stimulus evoked potentials. Stimulus evoked potentials for three different sensors. The large signal around time 0 is the stimulation artifact and the signals from 5-15 ms from stimulation onset are the evoked responses. Note the similarity between sEMG and Aichi DJ and the lack of sharp features in the OPM signals.

### Sensor noise floor required to capture voluntary MMG

We explored whether we could capture voluntary MMG with the Aichi sensors as well. To that end we used two sets of 4 Aichi DJ sensors aligned and stacked with 3 mm distance between each sensor. We then mounted the sensors on the extensor digitorum to attempt to capture similar signals across all sensors.

Figure 7a shows the SNR the highest SNR Aichi sensor during muscle contraction in a clench and hold task compared to periods of relaxation averaged across 50 trials. Applying ICA across all sensors resulted in a ∼0.1 increase in SNR across the bandwidth. Though not a significant enhancement, it nevertheless suggests we could leverage high density arrays with lower sensitivity to increase SNR when measuring voluntary MMG.

**Figure 7.**
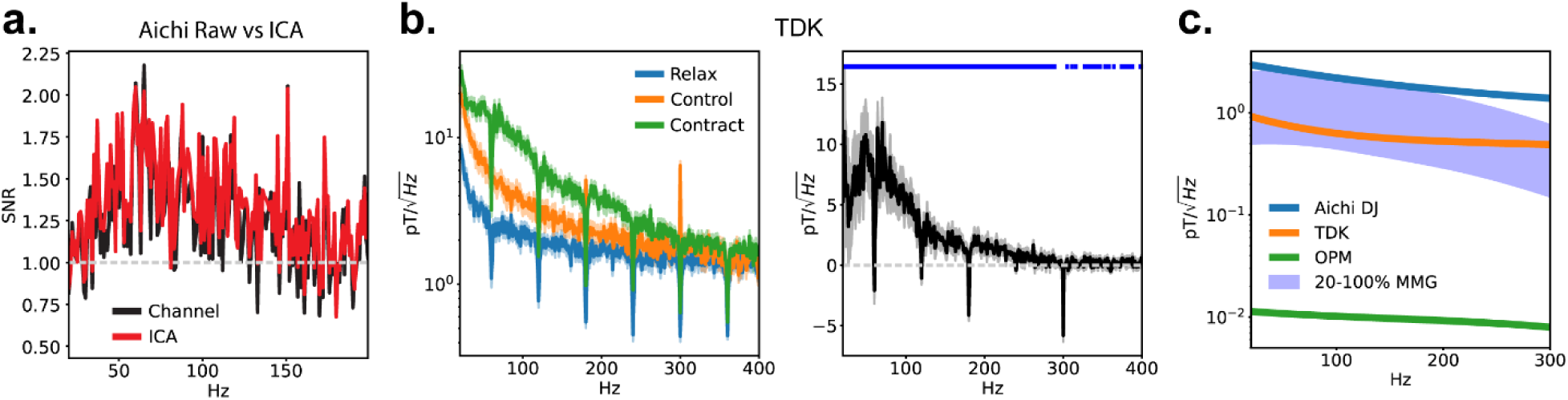
Voluntary MMG with non-OPM sensors. **a,** SNR of MMG captured with Aichi sensors and the ICA component with maximum SNR. **b,** MMG from voluntary movement captured with a TDK Nivio sensor. Averaged spectra during contraction (green), control sensor for movement artifact during contraction (orange), and during relax periods (blue) (left). Spectra during contraction subtracted by spectra of the control signal (right). Blue dots show statistical significance above zero (Wilcoxon rank sum test). **c,** Spectra of forearm MMG at 20-100% MVC when measured at the skin compared to the noise floors of the QuSpin OPM, TDK Nivio, and Aichi DJ. All spectra were estimated from measured values. Note the MMG power was calculated using signals captured by the OPMs with a correction on the higher frequency components to account for the sensor frequency response (Supplementary Figure 3).

We repeated the procedure with the TDK Nivio sensor (Figure 7b). The Nivio sensor was particularly sensitive to movement and it was especially challenging to securely fasten to the arm. Thus, we used another Nivio sensor with its sensitive axis down the length of the arm to purposefully pick up minimal MMG and used its measurements to estimate movement artifacts (Supplementary Figure 6). Averaging across 50 trials shows significant signals during contraction compared to the movement control up through 300 Hz.

These results suggest that the Aichi DJ sensor noise floor is at the threshold of measuring MVC MMG (Figure 7c). Though single TDK Nivio sensors were able to reliably capture large MMG signals, they struggled to discern signals during small forces of contraction. Though OPMs were able to fully measure MMG, we could not determine the change in signal strength at frequencies higher than 300 Hz. Nevertheless, our comparisons show that a sensor with a noise floor an order of magnitude higher than OPMs should be able to reliably capture MMG in this bandwidth.

## Discussion

### MMG reflects muscle activity similar to sEMG

Although magnetic signals recorded in the vicinity of skeletal muscle have been previously reported (Cohen & Givler, 1972; Zuo et al., 2020), there has yet to be validation that these signals directly reflect muscle activity and are directly analogous to sEMG. Muscle fibers generate large currents, but the magnetic signal may be obfuscated due to the orientations of the fibers and the asynchronous timing of the motor units (Broser et al., 2021); they may also be affected by signals generated by nerve bundles or even blood flow (Phua & Lissorgues, 2009; Wijesinghe, 2010). In addition, movement of the sensors, whether from muscle contraction, heart beats, or spontaneous tremors, could introduce artifacts if there is a residual magnetic field gradient present.

By measuring MMG simultaneously with sEMG at the same area we found that MMG shows very similar characteristics in spectral content, having a single peak at roughly the same frequency as well as similar bandwidth. In addition, we observed MMG to be highly linearly correlated with induced force, demonstrating it to be a direct measure of muscle contraction. We also found MMG can detect muscle fatigue at least as consistently as sEMG by comparing spectral content, demonstrating independent narrow band signals. MMG can thus be employed in applications requiring detection of degree of muscle contraction and small changes in signal properties, such as rehabilitation, athletics, or fine motor human-computer interfaces.

### MMG has distinct differences from sEMG

Though MMG displayed many similar properties to sEMG, we noticed a large difference between the high signal power content when comparing the SNR of the two modalities relative to frequency. Similarly, we also observed a larger change in median and mean frequencies in MMG compared to sEMG during fatigue suggesting more information content in high frequencies of MMG.

Spectral content in sEMG is heavily dependent on both the electrode size and the inter-electrode (IED) distance, with smaller sizes and shorter distances allowing for detection of higher frequencies (Afsharipour et al., 2019). As a result, although SENIAM (Surface ElectroMyoGraphy for the Non-Invasive Assessment of Muscles) recommended ∼10 mm size electrodes and ∼20 mm IED for bipolar sEMG recordings (Hermens et al., 2000), subsequent studies have suggested smaller electrodes with shorter IEDs to capture a larger bandwidth of the sEMG signal (Afsharipour et al., 2019; Alemu et al., 2003; Melaku et al., 2001).

We were limited to using the sEMG electrodes and distancing used in the study as we required non-ferrous contacts to prevent the OPMs from saturation or contaminate other sensor signals with noise. However, changing the IED from 20 mm to 10 mm causes the median frequency, or the frequency at which the total spectral power is split in half, to change only by roughly 5 Hz within a 500 Hz bandwidth (Rodriguez-Falces et al., 2015). Another study measuring mean frequency, or the average frequency weighted by the spectral power, corroborates these results showing a change of ∼3 Hz at IED of 18 mm compared to 36 mm (Alemu et al., 2003). As such, the IED alone does not fully explain the difference in the higher frequency power observed between MMG and sEMG.

The size of the electrodes potentially has a much larger effect on sEMG bandwidth as the increased area results in integration resulting in attenuation of higher frequencies (Merletti & Farina, 2016). However, previous studies have shown that for 10 mm wide devices the transfer function of spatial filtering results in -3 dB at 50 cycles / meter which translates to 200 cycles / second (Hz) when assuming 4 meters / second conduction velocity of muscle fiber signals (Afsharipour et al., 2019; Merletti & Muceli, 2019). Applying corrections based on these transfer functions as well as the OPM sensor’s frequency response showed that MMG still maintained significantly larger high frequency power (Supplementary Figure 4).

The difference in high frequency content is likely due to volume conduction affecting electrical signal transmission. Though MMG can be affected by passive currents traveling through tissue, the effect is minimal compared to the distortion caused by the changes in impedance at tissue boundaries (Hansen et al., 2010). We observed additional evidence of the spatial specificity of MMG compared to sEMG when comparing the minimum contraction they can detect, as MMG seemed to be much more sensitive to sensor location (Supplementary Figure 2). These results strongly suggests that MMG may provide better localization and could capture additional complex information compared to sEMG. It also alludes to possible use cases of MMG in applications requiring high frequency content, such as detection of muscle fatigue or assessment of motor unit action potentials.

One other large difference between MMG and sEMG is that MMG can operate with the same fidelity regardless of skin contact. However, as magnetic fields decrease in power with respect to distance, how much this advantage could be leveraged before the signals become too small to detect has not been fully quantified. Conflicting modeling results on whether the signal drops faster or slower with distance has caused further uncertainty (Parker & Wikswo, 1997; Wijesinghe, 1991). In this study we show conclusively that the signal from the surface of the skin drops slower than the inverse square law assuming the “center” of the signal source is less than 10 mm from the surface of the skin. We were still able to measure the MMG signal at greater than 50 mm from the surface of the skin, showing the non-contact aspect of MMG is indeed an advantage that can be utilized.

### Sensor technology is a major limiting factor

OPMs have been a major advancement in magnetic sensor development for biomagnetism. Compared to SQUIDs (superconducting quantum interference device), the original magnetometers used to record biomagnetism, they have comparable sensitivity while allowing for flexible positioning and high density, and also being significantly lower in cost. However, compared to sEMG sensors OPMs have lower bandwidth, smaller dynamic range, and are significantly more expensive.

The largest limitation in applicability of MMG is the lack of an appropriate sensor. For a magnetic sensor to enable MMG, it requires 1) high bandwidth, 2) large enough dynamic range to operate in unshielded conditions near electronic equipment and other magnetically noisy objects, 3) low noise floor to detect the relatively small MMG signals, 4) small size to allow for high density, and 5) low price for ubiquity. Although sensor development is still in progress, only three of the five constraints have been fulfilled by any single sensor. Though MMG may provide signals comparable to sEMG with potential advantages, a novel sensor technology is necessary to allow MMG to be readily employed in muscle sensing applications.

### Future directions

We have demonstrated the similarities of MMG to sEMG and highlighted some differences which may allow for better localization using MMG compared to sEMG. Though advantages in localization have been shown in models (Arekhloo et al., 2022; Klotz et al., 2022, 2023), there has yet to be empirical evidence. Magnetic sensing of the brain (magnetoencephalography (MEG)) has been demonstrated to provide higher localization accuracy compared to electrical sensing (electroencephalography (EEG)) in both a phantom as well as an implanted dipole in humans (Cohen et al., 1990; Cohen & Cuffin, 1991; Leahy et al., 1998), but the results cannot be extended to muscle sensing. The head has complex conductivity due to the thickness of the skull and boundaries between various tissues; in addition, brain sources are typically assumed to behave like dipoles (Hansen et al., 2010). In contrast, skeletal muscle is much closer to the surface of the skin and the fields produced by muscle fibers behave more similarly to finite wires (Parker & Wikswo, 1997; Wijesinghe, 1991). As such, there is a need to empirically compare localization performance between MMG and sEMG before drawing conclusions on their efficacies.

Additionally, though we were limited to the sEMG electrodes in our current study to prevent the OPM sensors from saturating, there is a wide range of sEMG sensors with highly varying properties, such as capacitive sensors and high-density EMG (Merletti et al., 2008; Sun & Yu, 2016). Though we estimated the effect of the electrode size on the spectral power, a more direct comparison will clarify the potential advantages of MMG.

Finally, we used OPMs in this study which allowed flexible and close placement of the sensors to the muscles but had clear limitations in bandwidth. Fortunately, there is ongoing research towards developing novel magnetometers to rival the sensitivity of OPMs while eliminating its disadvantages (Labanowski et al., 2016; Zuo et al., 2020). A magnetic sensor with higher bandwidth would allow for detection of voluntarily activated motor unit action potentials, potentially proving further insights into applications of MMG.

## Acknowledgements

We would like to thank Magnetic Shield Corporation for providing support in assessments of magnetic shielding, QuSpin for assistance in debugging and repairing the OPM sensors, Aichi Steel Corporation for granting details into their sensor electronics for customization, and TDK Corporation for the partnership in the Nivio sensor evaluation.

Additionally, we would like to thank the participants of the study for volunteering their time and permitting the use of their data.

## Author Contributions

R.Y., G.G., and I.G. conceived and designed the experiments. R.Y., G.G., I.G., R.C., D.D., and E.K. collected the data. R.Y., G.G., I.G., and R.C. performed analyses. R.Y. led drafting the manuscript and designing figures. All authors provided feedback on interpretation of the results and contributed to the final manuscript.

## Supplementary Figures

**Supplementary Figure 1.**
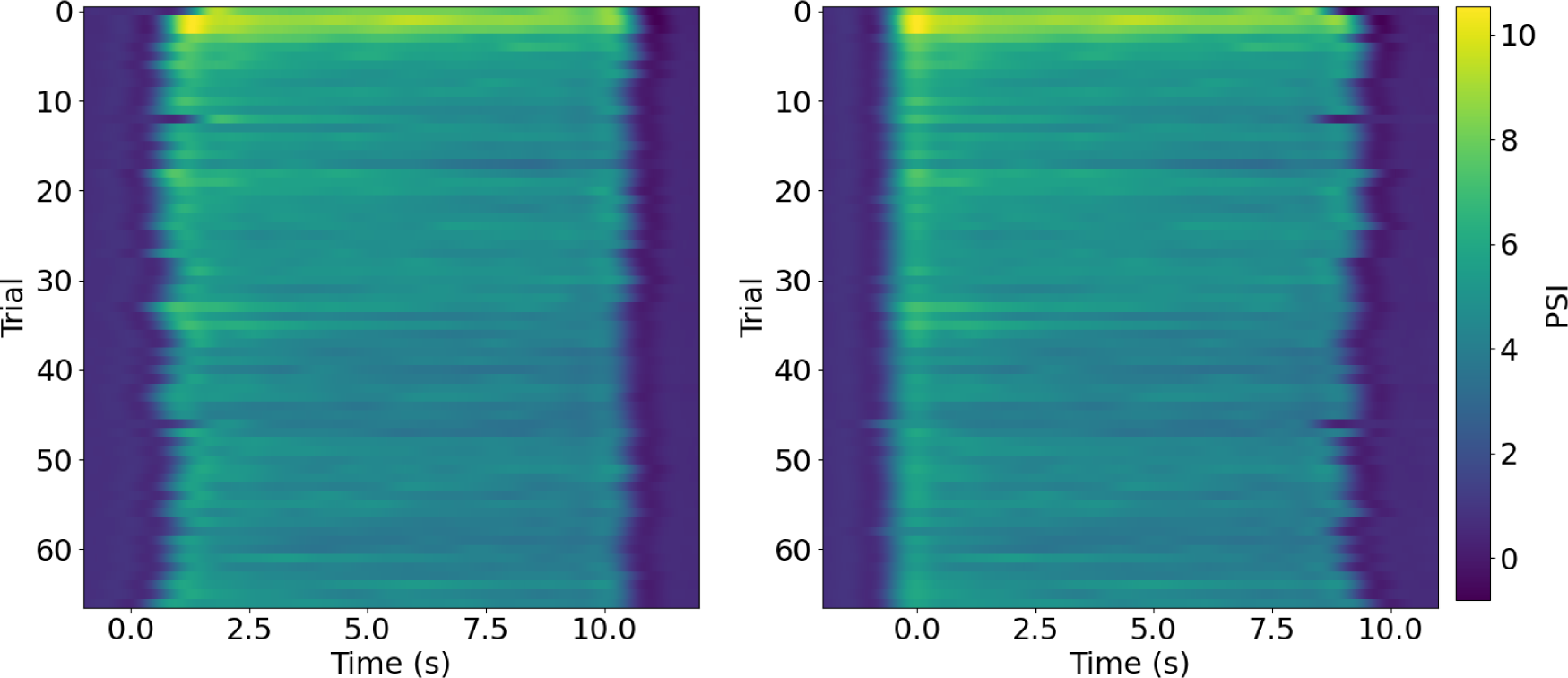
Alignment of trials using measured force. Subjects had changing reaction times to the tone to start each trial that widely varied over time and across subjects (left). Aligning each trial to the maximum contraction strength shows properly aligned trials allowing for consistent data analysis (right).

**Supplementary Figure 2.**
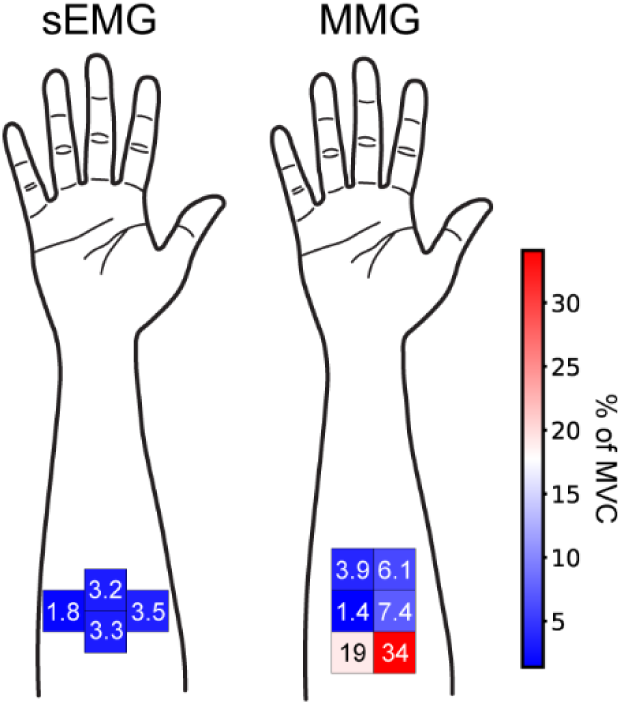
Minimum contraction strength detected by sEMG and MMG sensors. We calculated the minimum contraction strength a sensor can detect by the degree of force required to reach the baseline amplitude plus two times the standard deviation of the baseline. Note that MMG was as sensitive to low contraction forces as sEMG but seemed to also be more reliant on the specific positioning of the sensor.

**Supplementary Figure 3.**
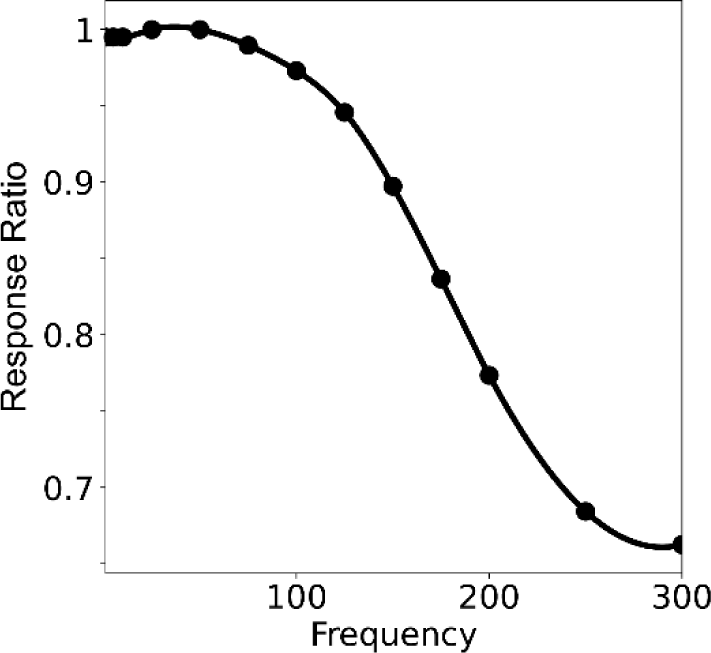
Frequency response of QuSpin OPM sensors. Frequency response was measured by playing a magnetic tone at various frequencies (dots) at the same amplitude. The curve across the bandwidth was obtained using cubic interpolation.

**Supplementary Figure 4.**
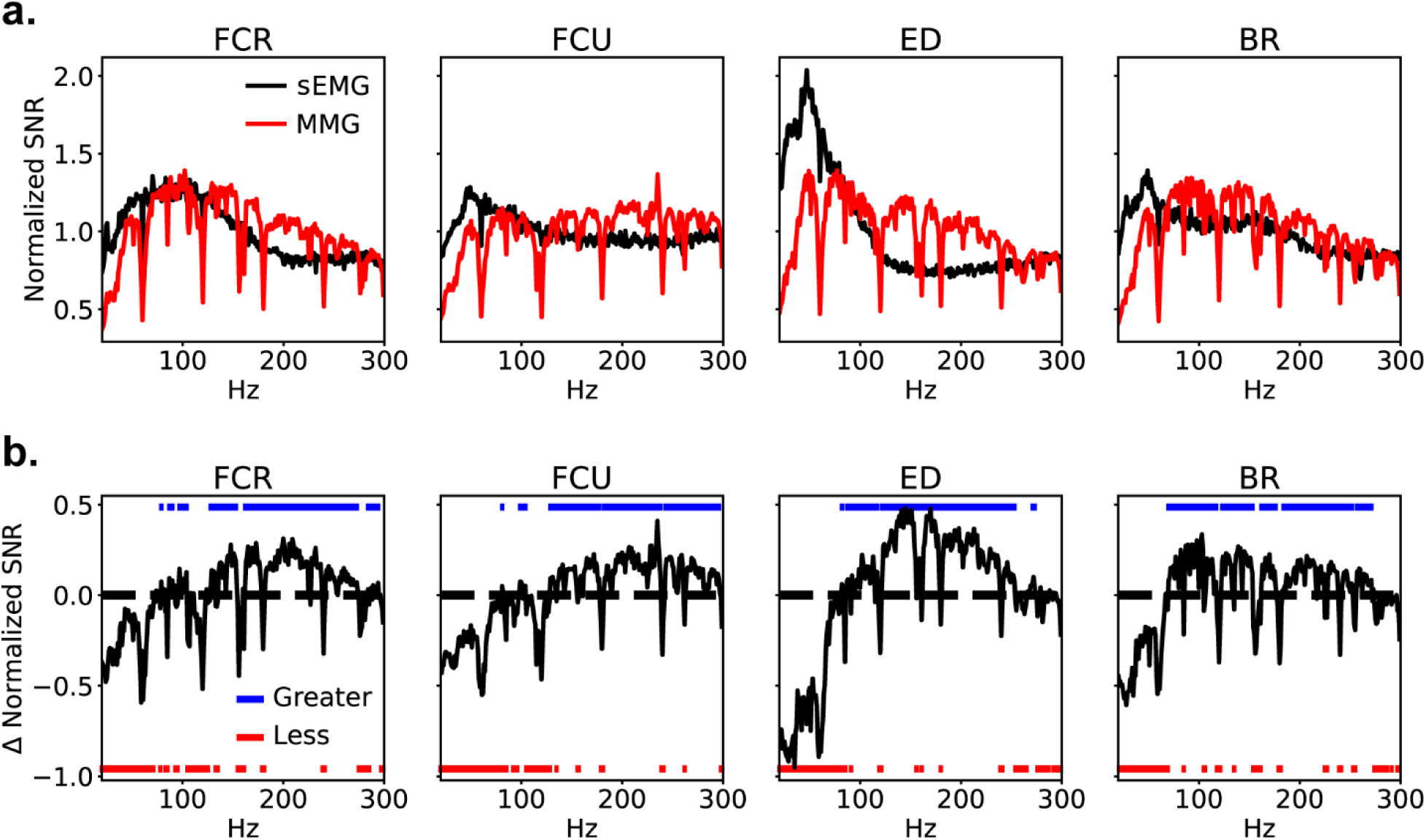
Frequency analysis with MMG and sEMG corrected with sensor frequency responses. Analysis analogous to Figure 3. **a,** Normalized SNR of sEMG and MMG and **b,** the difference between MMG SNR and sEMG SNR. The MMG SNR was corrected by using the frequency response of the OPM sensors (Supplementary Figure 4) and the sEMG SNR was corrected to match an electrode with 3 mm diameter using an estimated frequency response.

**Supplementary Figure 5.**
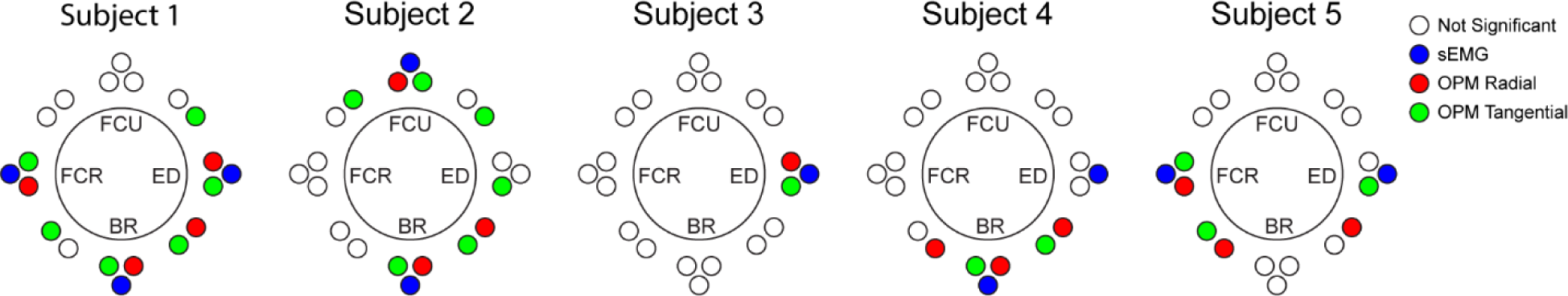
Fatigue per subject. Channels that showed signs of fatigue per subject both within and across trials, see Figure 1 for sensor placements. Significance was determined by comparing mean and median frequencies of the first 1 second or 10 trials to the last 1 second or 10 trials to test for changes both within and across trials. Fatigue was only noted if the differences were statistically significant (Wilcoxon rank-sum test) for both comparisons. Note both sEMG and MMG (OPMs) show similar locations of fatigue per subject.

**Supplementary Figure 6.**
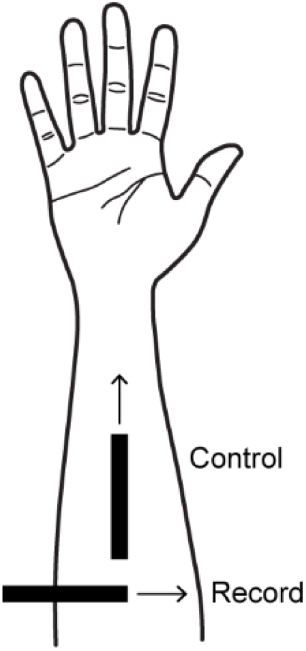
Positioning of TDK sensors. Placement of TDK sensors (black bars) on the forearm with sensitive axes (arrow) labeled for each sensor. The control sensor picks up minimal MMG as the sensitive axis is along the length of the muscle fibers.

